# Reliability of single-subject neural activation patterns in speech production tasks

**DOI:** 10.1101/807925

**Authors:** Saul A. Frankford, Alfonso Nieto-Castañón, Jason A. Tourville, Frank H. Guenther

**Author notes:** Email addresses (S.A. Frankford); (A. Nieto-Castañón); (J.A. Tourville); (F.H. Guenther). Send correspondence to: Saul A. Frankford, Boston University, Department of Speech, Language and Hearing Sciences, 677 Beacon St., Boston, MA 02215.

## Abstract

Speech neuroimaging research targeting individual speakers could help elucidate differences that may be crucial to understanding speech disorders. However, this research necessitates reliable brain activation across multiple speech production sessions. In the present study, we evaluated the reliability of speech-related brain activity measured by functional magnetic resonance imaging data from twenty neuro-typical subjects who participated in two experiments involving reading aloud simple speech stimuli. Using traditional methods like the Dice and intraclass correlation coefficients, we found that most individuals displayed moderate to high reliability. We also found that a novel machine-learning subject classifier could identify these individuals by their speech activation patterns with 97% accuracy from among a dataset of seventy-five subjects. These results suggest that single-subject speech research would yield valid results and that investigations into the reliability of speech activation in people with speech disorders are warranted.

## 1. Introduction

Our understanding of the neural mechanisms responsible for speech and language has dramatically improved in recent decades due to the development of non-invasive techniques for measuring whole-brain activity. Perhaps the most widely used technique of this type is functional magnetic resonance imaging (fMRI); at least 4,500 papers have been published on this topic in pubmed since 2000^1^. To date, the vast majority of fMRI studies of speech and language have involved analyzing group average results from cohorts of 10 or more neurotypical participants, in many cases compared to similar-sized cohorts of patients with neurological conditions that impact speech or language function. Collectively, these studies have revealed a network of brain areas that are commonly active during speech production (Guenther, 2016; Price, 2012). When brain responses are compared between groups, however, the results are often less consistent (e.g., Connelly et al., 2018 vs. Chang et al., 2009). This could result from the relatively small sample sizes of typical fMRI study designs lacking sufficient power, a shortcoming that is being addressed in more recent studies with larger samples sizes and data pooling (Brown et al., 2005; Costafreda, 2009; Turkeltaub et al., 2002), i.e., measuring across larger groups.

Larger groups, however, cannot address another factor that is becoming more apparent to those mapping the functional components of the speech production network: high between-subjects variability in the location and level of speech-related BOLD responses. Attempts to localize the locus of “crucial” neural damage in acquired apraxia of speech (AOS), for instance, have reported a variety of locations (Dronkers, 1996; Hillis et al., 2004; Moser et al., 2016). Moreover, there is tremendous variability in the location and extent of stroke-related damage to neural tissue across *individuals*. This individual variability found in AOS and other speech network disturbances (e.g., stuttering, Wymbs et al., 2013) can mask group differences in fMRI analyses, and make it difficult to map the neural locus (or loci) of a given disorder.

An alternative approach for studying speech disorders is to use subject-specific study designs that are unaffected by between-subjects variability. A number of studies have demonstrated the utility of single-subject fMRI study designs or encouraged its future use for a range of purposes. These include mapping language areas prior to resective surgery for patients with epilepsy or gliomas (Babajani-Feremi et al., 2016; Bizzi et al., 2008; Chen & Small, 2007; Gross & Binder, 2014) improving diagnosis of disorders (Raschle et al., 2012; Sundermann et al., 2014), and determining whether neural plasticity following stroke can predict outcomes (Chen & Small, 2007; Kiran et al., 2013; Meltzer et al., 2009). In the speech domain, single-subject approaches have been used to evaluate responses to treatment in AOS (e.g., Farias et al., 2014), but these could be expanded to tracking natural neural organization changes over time in developmental speech disorders like stuttering. Due to the individuality of the presentation of these disorders, subject-specific approaches could provide more meaningful measures of change not captured in group average analyses.

However, the suitability of subject-specific studies of speech and language processes, depends heavily on the reliability of speech-related activity in individual brains. The main purpose of the current study is to test this assumption by assessing the reliability of single-subject fMRI measured during speech production tasks across scanning sessions. Several prior studies have examined within-subject reliability of BOLD responses during language production tasks (e.g. Mayer, Xu, Paré-Blagoev, & Posse, 2006; Otzenberger, Gounot, Marrer, Namer, & Metz-Lutz, 2005; Wilson, Bautista, Yen, Lauderdale, & Eriksson, 2017). Many have used a covert speech task (Brannen et al., 2001; Harrington et al., 2006; Maldjian et al., 2002; Mayer et al., 2006; Otzenberger et al., 2005; Rutten et al., 2002) or have focused on a limited set of regions of interest (ROIs) like Broca’s area and temporo-parietal cortex (e.g., Brannen et al., 2001; Harrington et al., 2006; Mayer et al., 2006; Otzenberger et al., 2005; Rau et al., 2007). However, speech requires overt motor actions and the integration of sensory feedback supported by large and often distant areas of the brain (Guenther, 2016; Sato, Vilain, Lamalle, & Grabski, 2015). Four recent studies (Gorgolewski et al., 2013; Nettekoven et al., 2018; Paek et al., 2019; Wilson et al., 2017) have assessed reliability in neurologically normal participants across the cortex during overt word production. These studies report moderate to high levels of reliability and each provides unique insight into the factors that impact test-retest reliability, especially pertaining to older adults and clinical populations.

Our aim in the present study was to determine whether such reliability is robust in the speech production network across different speaking tasks and interscan intervals. To do this, we performed a retrospective analysis of participants who had taken part in more than one fMRI study of speech production in our lab. This had the advantage of assessing the reliability of general speech network activation patterns in an individual rather than the reliability of a specific task to allow for greater generalization of the results herein. Compared to previous work, we included studies with stimuli that limited higher-level linguistic processing. Doing so allowed us to assess the reliability of neural activity specific to speech motor control processes. Finally, since these datasets were collected for basic research purposes in healthy individuals, they were composed of much longer sessions which may improve the reliability of an individual’s speech network activity.

We used the Dice coefficient to measure the spatial overlap of active brain regions within individuals across multiple speech production studies. This easily interpretable measure can be compared to numerous previous studies of fMRI reliability (Bennett & Miller, 2010). For a more thorough reliability measure that accounts for both the location and relative scale of activity across the brain, we calculated a single-subject intraclass correlation coefficient (ICC; as in Raemaekers et al., 2007). While each of these provides an estimate of similarity that can be used in a single-subject context, further information can be gleaned from measures that assess reliability in relation to a between-subjects standard. We therefore computed an ICC for each vertex on the cortical surface to yield a map of reliability (as in Aron, Gluck, & Poldrack, 2006; Caceres, Hall, Zelaya, Williams, & Mehta, 2009; Freyer et al., 2009; Meltzer et al., 2009). This measure estimated the reliability and discriminability of activation across the entire brain at a vertex level. Finally, we directly tested whether an individual speaker’s neural activation patterns during speech in one study could predict activation in a second study using a machine learning classifier.

Reliability measures were compared to two benchmarks: a chance-level baseline derived from random data maps, and a residual signal map derived from anatomy-related information in the BOLD signal that we would expect to have high reliability.

## 2. Materials and Methods

### 2.1. Participants

Our dataset comprises seventy-five individuals who previously participated in fMRI studies of speech production in the SpeechLab at Boston University. Of these, data from twenty individuals (mean age: 28.95 years, range: 19-44, 10 female/10 male) who participated in at least two fMRI studies (see Tables 1 and 2) were used to evaluate reliability (median number of days between studies: 13.5, range: 6 - 196). Data from the remaining fifty-five speakers (age range: 18-51) from these or three other speech production studies (see Table 2) were added in the classifier analysis to train the subject classifier and to generalize its features to the broader population of healthy speakers (see section 2.5.4. Subject Classifier). All participants were right-handed native speakers of American English and reported normal or corrected-to-normal vision as well as no history of speech, language, hearing, or neurological disorders. Informed consent was obtained from all participants, and each study was approved by the Boston University Institutional Review Board.

**Table 1.**
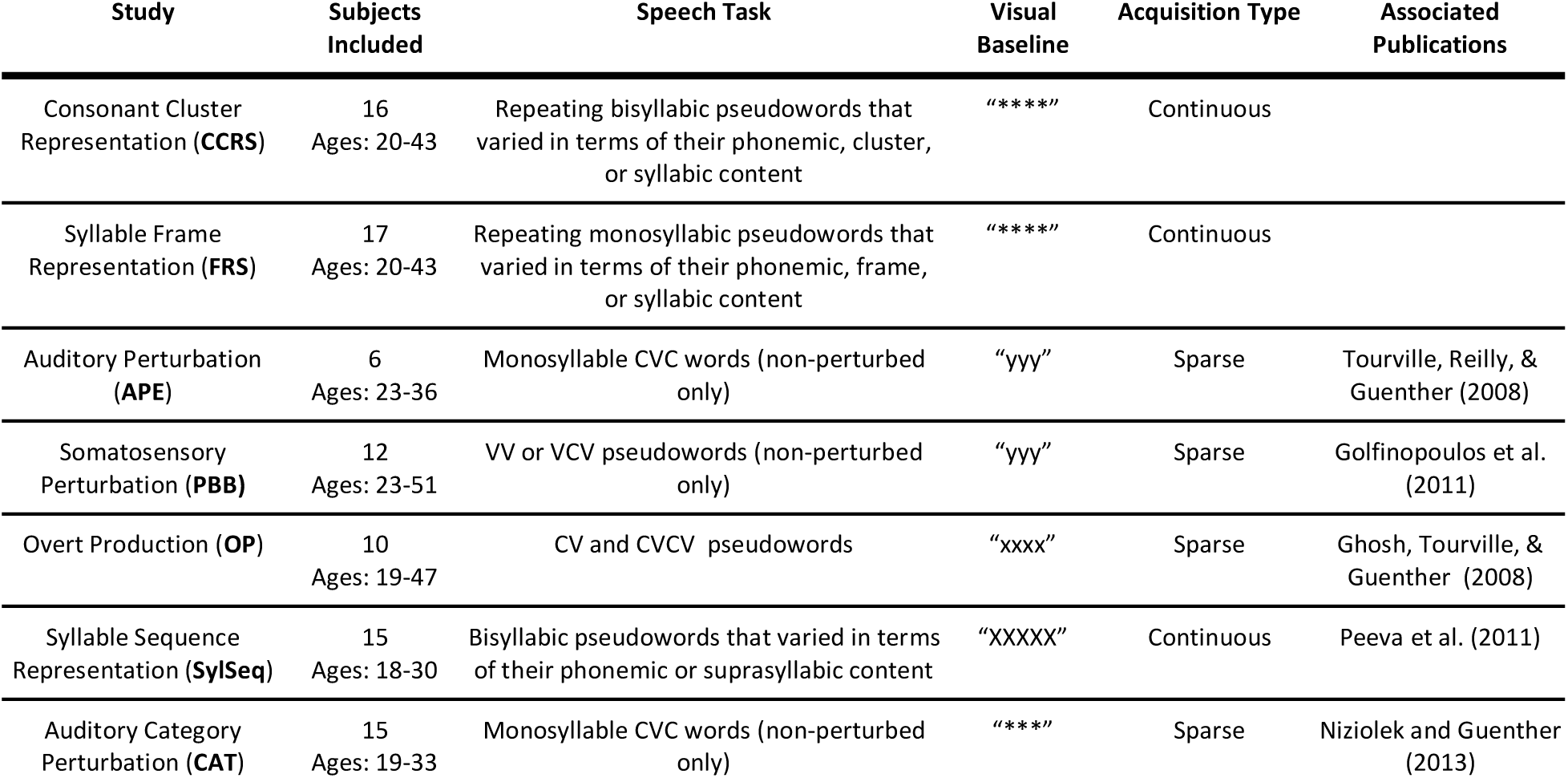
Information about the studies from which activation maps were included in the present analyses. C = consonant, V = vowel.

**Table 2.** Studies in which each test subject participated, total number of trials, and time between studies. Study identification codes refer to abbreviations in the ‘Study’ column of Table 1.

| Subject | Study 1 |  |  | Study 2 |  |  | Days Between Studies |
| --- | --- | --- | --- | --- | --- | --- | --- |
|  | ID | Speech Trials | Baseline Trials | ID | Speech Trials | Baseline Trials |  |
| 1 | CCRS | 378 | 126 | FRS | 258 | 66 | 6 |
| 2 | CCRS | 378 | 126 | FRS | 258 | 66 | 14 |
| 3 | CCRS | 324 | 108 | FRS | 258 | 66 | 52 |
| 4 | CCRS | 378 | 126 | FRS | 258 | 66 | 7 |
| 5 | CCRS | 324 | 108 | FRS | 258 | 66 | 6 |
| 6 | CCRS | 378 | 126 | FRS | 258 | 66 | 20 |
| 7 | CCRS | 324 | 108 | FRS | 258 | 66 | 7 |
| 8 | CCRS | 378 | 126 | FRS | 258 | 66 | 13 |
| 9 | CCRS | 378 | 126 | FRS | 258 | 66 | 19 |
| 10 | CCRS | 378 | 126 | FRS | 258 | 66 | 12 |
| 11 | CCRS | 324 | 108 | FRS | 258 | 66 | 7 |
| 12 | CCRS | 324 | 108 | FRS | 258 | 66 | 7 |
| 13 | CCRS | 378 | 126 | FRS | 258 | 66 | 7 |
| 14 | CCRS | 324 | 108 | FRS | 258 | 66 | 7 |
| 15 | APE | 191 | 64 | PBB | 192 | 32 | 75 |
| 16 | APE | 191 | 64 | PBB | 144 | 24 | 163 |
| 17 | APE | 191 | 64 | PBB | 192 | 32 | 196 |
| 18 | APE | 187 | 63 | PBB | 240 | 40 | 21 |
| 19 | APE | 192 | 64 | PBB | 192 | 32 | 70 |
| 20 | APE | 143 | 48 | PBB | 240 | 40 | 28 |

### 2.2. Speech Tasks

All speech tasks included in the present study were overt productions of either real words or pseudowords formed by two or more consecutive phonemes. These characteristics ensure a distribution of tasks used in neuroimaging studies of speech, while limiting activation patterns to those associated with overt speech production that includes phonemic transitions. A list of speaking tasks and their visual baseline control conditions from each study is included in Table 1. Details of the four studies from which repeated measures were taken (CCRS, FRS, APE, and PBB) are described here. More detailed information on the other studies (OP, SylSeq, and CAT) is provided in the publications listed in Table 1.

The CCRS and FRS experiments were block-design fMRI studies in which subjects produced sequences of pseudowords during continuous scanning. Both studies included multiple speech conditions and a baseline condition. During speech trials, subjects simultaneously viewed an orthographic representation and heard a recording of the pseudoword to be produced. A white cross replacing the orthographic representation cued the subject to produce the pseudoword. On baseline trials, subjects saw a series of asterisks on the screen rather than orthographic stimulus and rested quietly. Functional runs were organized into blocks of 6 trials of the same condition with a 3 s pause between blocks. Pseudowords and conditions were randomized within runs.

Sequences in the CCRS study comprised pairs of two-syllable pseudowords that varied in the number of unique phonemes, consonant clusters and syllables in the sequence. The conditions were: exact repetition (e.g., ‘GROI SLEE, GROI SLEE’); same phonemes and consonant clusters, different syllables (e.g. ‘GROI SLEE, GREE SLOI’); and different phonemes, consonant clusters, and syllables (e.g. ‘KWAI BLA, SMOO KROI’). Each trial lasted 2.5 s. Runs consisted of fifteen blocks, and lasted approximately 5 min. Each subject completed 7 runs that optimally allowed for approximately 21 blocks per condition per subject. In total, 120 fMRI volumes were acquired continuously during each run.

Sequences in the FRS study were pairs of monosyllabic pseudowords that varied in the number of unique phonemes, syllables, and syllabic frames (see MacNeilage, 1998). The conditions were: exact repetition (e.g. ‘TWAI, TWAI’); same frames, different phonemes and syllables (e.g. ‘FAS REEN’); same phonemes, different frames and syllables (e.g. ‘RAUD DRAU’); and different frames, phonemes, and syllables (e.g. ‘DEEF GLAI’). Each trial lasted 2 seconds. Runs consisted of eighteen blocks and lasted approximately 4.5 min. Each pseudoword or pseudoword pair was maximally used once per block and in 2-3 blocks throughout the experiment to maintain novelty. Each subject completed 6 runs that optimally allowed for approximately 27 blocks per condition per subject. In total, 108 fMRI volumes were acquired continuously during a run.

The APE (Tourville et al., 2008) and PBB studies (Golfinopoulos et al., 2011), used a sparse fMRI acquisition design that allowed subjects to produce speech during silent intervals between fMRI volume acquisitions. In both experiments, subjects were instructed to read aloud the speech stimulus presented orthographically at the onset of each trial or to remain silent if a control stimulus (the letter string ‘yyy’) was presented. Stimuli in the APE study consisted of 8 /CεC/ words (e.g., beck, bet, debt). Stimuli remained onscreen for 2 s. An experimental run consisted of 64 speech trials (8 presentations of each word) and 16 control trials (Tourville et al., 2008). On 25% of speech trials, the first formant (F1) of the subject’s speech was altered before being fed back to the subject. Trial order was randomly permuted within each run such that consecutive presentation of the same stimulus and consecutive F1 shifts in the same direction were prohibited. Subjects performed 3 or 4 functional runs. *Only speech trials with normal feedback* and baseline trials were included in the present study.

Speech stimuli in the PBB study (Golfinopoulos et al., 2011) consisted of eight pseudowords that required a jaw closure after producing an initial vowel (e.g., /au/, /ani/, /ati/). Stimuli remained onscreen for 3 s. Each experimental run consisted of 56 speech trials (seven presentations of each pseudoword) and 16 baseline trials. On one seventh of all speech trials and half of all baseline trials, jaw closure was restricted by the rapid inflation of a small balloon positioned between the subjects’ upper and lower molars. Trial order was randomly permuted within each run such that consecutive perturbation trials were prohibited. Subjects included in the present analysis completed between three and five runs. *No perturbation trials* were included in the present analysis.

### 2.3. Image Acquisition

MRI data were acquired at the Athinoula A. Martinos Center for Biomedical Imaging at Massachusetts General Hospital (APE, PBB, OP, CCRS, FRS), the Athinoula A. Martinos Imaging Center at the McGovern Institute for Brain Research at the Massachusetts Institute of Technology (CAT), and the fMRI Centre of Marseille (SylSeq).

For CCRS and FRS, data were acquired using a 3 Tesla Siemens Trio Tim scanner with a 32-channel head coil. For each subject, a whole-brain high-resolution T1-weighted MPRAGE volume was acquired (voxel size: 1 mm^3^, 256 sagittal images, TR: 2530 ms, TE: 3.44 ms). T2*-weighted volumes consisting of 41 gradient echo – echo planar axial images (in plane resolution: 3.1 mm, slice thickness: 3 mm, gap: 25%, TR: 2.5 s, TA: 2.5 s, TE: 20 ms) were collected continuously during functional runs.

For APE and PBB, a high-resolution T1-weighted anatomical volume (128 slices in the sagittal plane, slice thickness: 1.33 mm, in-plane resolution: 1 mm^2^, TR: 2530 ms, TE: 3.3 ms) was obtained for each subject prior to functional imaging. Functional volumes consisted of 32 gradient echo - echo planar axial images (in plane resolution: 3.125 mm^2^, slice thickness: 5 mm, TR: 2000 ms, TE: 30 ms). A sparse sampling (Hall et al., 1999) clustered volume acquisition method, consisting of silent intervals between consecutive volume acquisitions, was used. Two consecutive volumes (each volume acquisition taking 2 s) were acquired 5 s after the onset of each trial.

See Peeva et al. (2010), Ghosh, Tourville, & Guenther (2008), and Niziolek & Guenther (2013) for acquisition parameters for the SylSeq, OP, and CAT studies, respectively (refer to Table 1 for study codes).

### 2.4. Preprocessing and first-level analysis

Preprocessing was carried out using SPM12 (http://www.fil.ion.ucl.ac.uk/spm) and the CONN toolbox (Whitfield-Gabrieli & Nieto-Castanon, 2012) preprocessing modules. Each participant’s functional data were motion-corrected to their first functional image, and coregistered to their structural image using SPM12’s inter-modality coregistration procedure with a normalized mutual information cost function (Collignon et al., 1995; Studholme et al., 1998). For CCRS and FRS, BOLD responses were high-pass filtered with a 128-second cutoff period and estimated at each voxel using a general linear model (GLM). The hemodynamic response function (HRF) for each stimulus block was modeled using a canonical HRF convolved with the trial duration from each study. For APE and PBB, the BOLD response for each event was modeled using a single-bin finite impulse response (FIR) basis function spanning the time of acquisition of the two consecutive volumes. For each run, a linear regressor was added to the model to remove linear effects of time, as were six motion covariates and a constant session effect (the intercept for that run). See Peeva et al. (2010), Ghosh, Tourville, & Guenther (2008), and Niziolek & Guenther (2013) for first-level design details in the other studies. Functional data were also censored (Power et al., 2014) by including additional regressors for all studies to remove the effects of volumes with excessive motion and global signal change, as identified using ART (https://www.nitrc.org/projects/artifact_detect/) with a scan-to-scan motion threshold of 0.9 mm and a scan-to-scan signal intensity threshold of 5 standard deviations above the mean.

In all studies and subjects, first-level model estimates for each speech condition and baseline were contrasted at each voxel and averaged across all study-specific speech conditions to obtain speech activation maps (*speech* maps). Effect size maps were used for subsequent analyses rather than significance (*p*-value) maps because a) significance maps are not as consistent for individual subjects as they are for group analyses (Gross & Binder, 2014; Voyvodic, 2012) and b) previous research has demonstrated greater overlap in effect size maps (Wilson et al., 2017). T1 volume segmentation and surface reconstruction were carried out using the FreeSurfer image analysis suite (freesurfer.net; Fischl, Sereno, & Dale, 1999). Activation maps were then projected to each individual’s inflated structural surface. To align subject data, individual surfaces were inflated to a sphere and coregistered with the FreeSurfer mean surface template (fsaverage; see Figure 1). Surface maps were then smoothed using iterative diffusion smoothing with 40 diffusion steps (equivalent to a 8 mm full-width half maximum smoothing kernel, Hagler et al., 2006). This level of smoothing has previously been shown to optimize reliability of task-related BOLD response data in individuals (Caceres et al., 2009).

**Figure 1.**
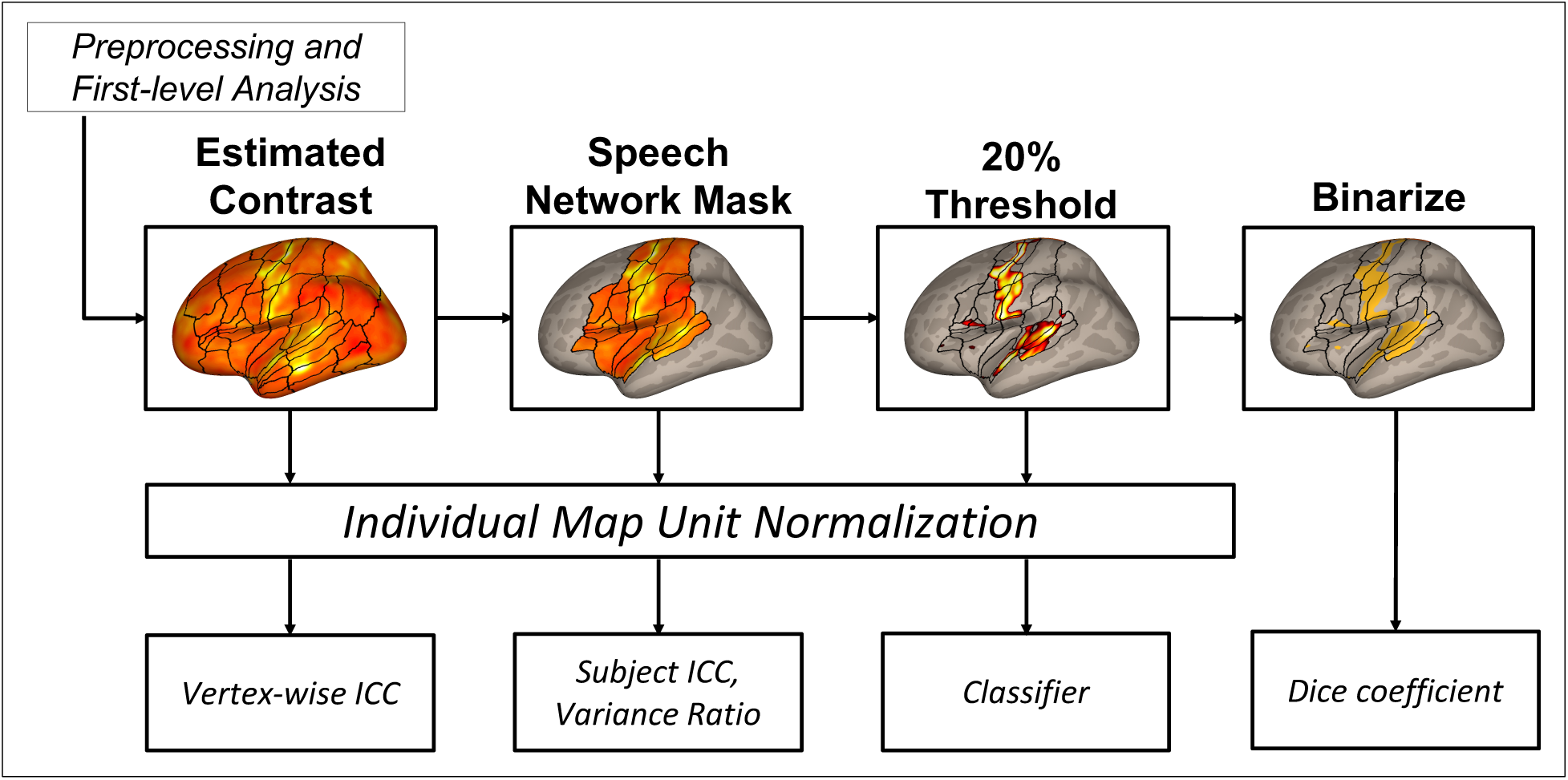
Thresholding pipeline map for each of the reliability analyses. After preprocessing and estimation of first-level condition effects, the *speech, null*, and *random* maps were calculated, and submitted to the vertex-wise ICC analysis. A speech network mask was applied, so that only vertices inside this mask were used for the single-subject ICC and variance ratio measures. Next, the 20% of vertices with the highest activation levels were kept for the classifier analysis. Finally, these thresholded maps were binarized for the Dice coefficient analysis. Prior to calculating reliability measures (except the Dice coefficient), maps were normalized to account for differences in effect size scaling between subjects and studies. Outlines for regions of interest previously described in Tourville & Guenther (2012) are included for reference, and appear only in areas of cortex on which a given analysis was carried out.

In addition to the above *speech* maps, we computed two other sets of maps for comparison purposes. The first was *random* maps, representing randomly generated data with similar spatial properties, and processed in exactly the same way as the *speech* maps. We expected these maps to show minimal reliability (chance-level). Reliability measures derived from *random* maps served as a baseline reference, and to eliminate the possibility that our preprocessing and estimation procedure would artifactually introduce unexpected biases in reliability metrics. The second was *null* maps, representing anatomical information about each subject like tissue morphology and neurovasculature present in the average BOLD signal, and, again, were processed in exactly the same way as the *speech* maps. We expected these maps to show high reliability, as anatomical information is expected to vary minimally over the time spans considered in this study. Reliability measures derived from *null* maps served as references for comparison purposes, and to explore the possibility that reliability of speech-related functional activation may be influenced by, or related to, reliability of anatomical features.

Maps of random activation (*random* maps) were created by independently replacing effect sizes at each vertex with a randomly chosen value from a normal distribution (mean of 0 and a standard deviation of 1) and smoothing the data to the same degree as the *speech* maps. To obtain maps of average MRI signal (*null* maps) that is not affected by task effects, estimates of the constant regression term of each run were averaged for each subject in each study. These maps represent the average T2* signal after the effects of speech, baseline, motion, and outliers have been removed. Similar to the *speech* maps, they were then projected to each individual’s structural surface. Because there is individual variability in the T2* signal across the cortex, these maps represent individual features of a subject’s cortical anatomy

### 2.5. Reliability Measures

We used two measures to quantify individual-subject activation reliability across different sessions in individuals (while sessions come from two separate studies, for clarity the term *session* will be used going forward to refer to a data collection time point): the Dice coefficient and a single-subject intraclass correlation coefficient. Two further measures were used to examine sampled-normed reliability: a vertex-wise intraclass correlation coefficient, and a machine-learning classifier. Each of these measures was applied to the *speech, random*, and *null* maps.

#### 2.5.1. Single-subject Spatial Overlap

To measure the spatial overlap of supra-threshold vertices, we used the Dice coefficient, a metric widely used in fMRI reliability studies (see Bennett & Miller, 2010 for a review). It is the ratio between the extent of overlap of individual maps and their average size and yields values between 0 (no overlap) and 1 (complete overlap). A strength of this measure is that it is straightforward to interpret and provides a simple way to characterize the reproducibility of thresholded activation maps (Bennett & Miller, 2013). On the other hand, the Dice coefficient is sensitive to how these maps are thresholded (Duncan et al., 2009; Smith et al., 2005), and the area over which the calculation is made (Gorgolewski et al., 2013), where lower thresholds and whole-brain analyses will tend to increase overlap. Despite this, the Dice coefficient provides a rough estimate of neural response reliability.

The Dice coefficient is formally given by:

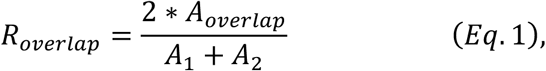

where *A*_*1*_ and *A*_*2*_ are defined as the number of supra-threshold vertices for individual sessions and *A*_*overlap*_ is the total number of vertices that exceeds the threshold in both sessions (Bennett & Miller, 2010). Because we were only interested in assessing reliability in brain areas commonly activated during speech production, we masked each map to only analyze activation within a predefined speech production network area covering approximately 35% of cortex (see Figure 1; Tourville & Guenther, 2012). Activation maps were then thresholded to retain only the highest 20% of surface vertices within the masked area (approximately 7% of total cortex; see Figure 2 for examples of these thresholded maps). Finally, this map was binarized (active voxels = 1, all other voxels = 0).

**Figure 2.**
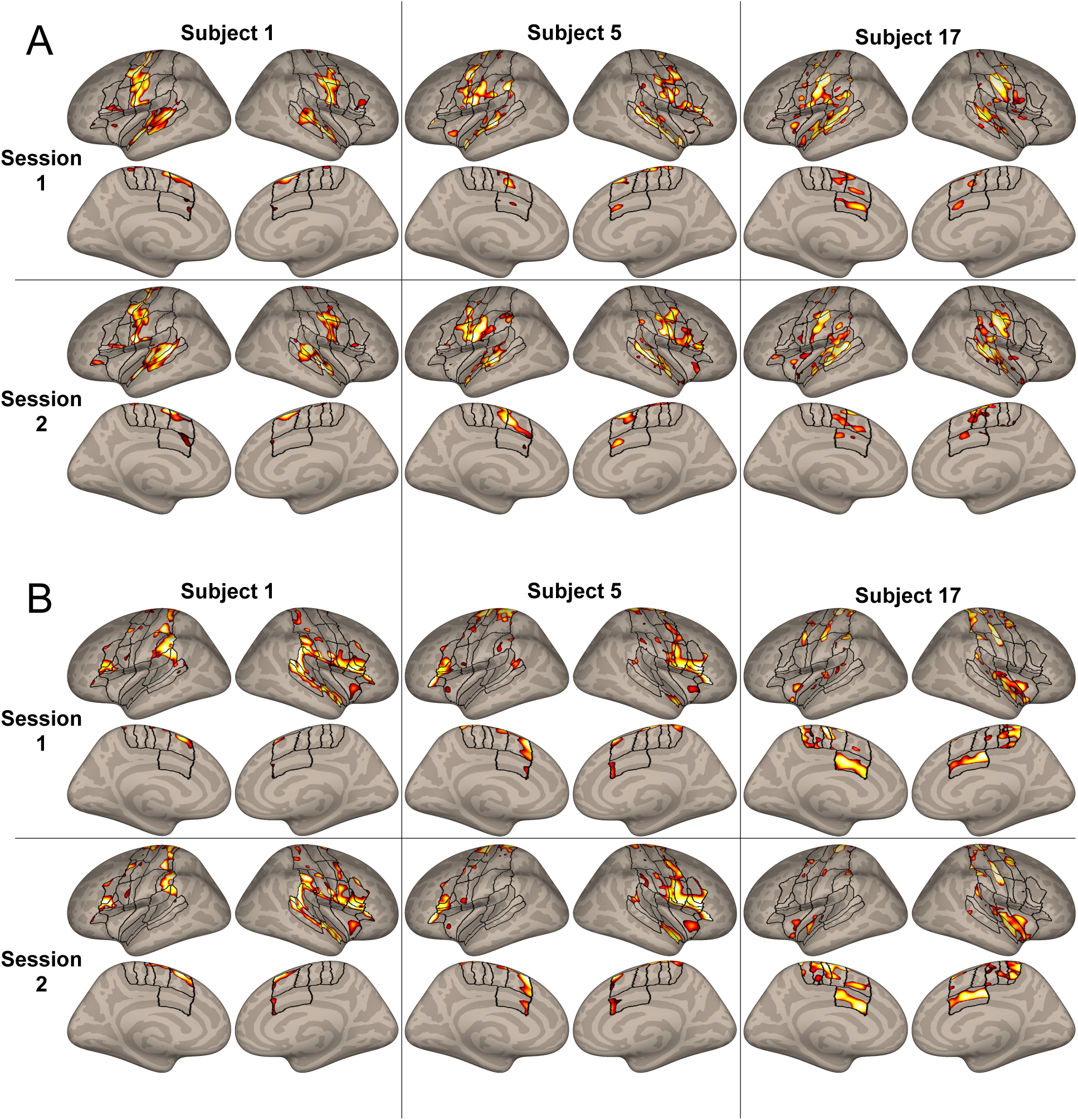
A. Masked and thresholded *speech* maps for three example subjects in both sessions. Outlines of regions of interest covering the masked speech network previously described in Tourville & Guenther (2012) are included for reference. B. Masked and thresholded *null* maps for the same subjects. In both cases, the activation peaks display broad visual similarity between sessions. Note: the color scale indicates the rank of vertex activation within each map, where lighter colors indicate higher activation.

#### 2.5.2. Single-subject ICC

To obtain a measure of reliability that was not threshold-dependent and took into account the level of activation at each vertex, we calculated a single-subject ICC (see Raemaekers et al., 2007) for each subject that compares variance between sessions to within-session (across-vertex) variance. Like the Dice coefficient, the ICC is relatively straightforward to interpret: a value of 0 means there is no correlation across all vertices, while a value of 1 signifies perfect correlation across all vertices. Of the many types of ICCs described in the literature, we used the ICC(1) as defined in McGraw and Wong (1996). This type of ICC is based on an analysis of variance (ANOVA) of the following one-way random effects model:

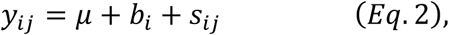

where *y*_*ij*_ is the value for the *i*^*th*^ vertex and the *j*^*th*^ session, *μ* is the mean value across all vertices and sessions, *b*_*i*_ is the between-vertices effect at vertex *i*, and *s*_*ij*_ is the residual, representing the between-sessions effect. ICC(1) estimates the degree of absolute agreement across multiple repetitions of a set of measurements. Formally, it is an estimate of

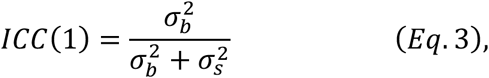

where 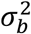 is the between-vertex variance and 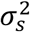 is the between-sessions variance. Based on McGraw and Wong (1996), the sample estimate, 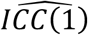, can be calculated using the following formula:

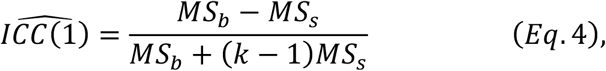

where *MS*_*b*_ is the mean squares across vertices, *MS*_*s*_ is the mean squares of the residuals, and *k* is the number of within-subjects measurements (in this case, 2 sessions). Following the convention of Koo and Li (2016), ICC values below 0.5 indicate poor reliability, between 0.5 and 0.75, moderate reliability, between 0.75 and 0.9, good reliability, and above 0.9, excellent reliability.

In addition, to determine whether reliability in individual subjects across sessions was higher than that across the sample, we also computed a between-subjects ICC analysis. This was accomplished by averaging each individual’s *speech* maps across sessions, and estimating the same ICC defined in *Eq*. 2 and *Eq*. 3. Thus, the *s* term estimated the between-subjects effect rather than the between-session effect.

For this analysis, activation maps were masked with the same speech production network mask described for the overlap analysis but no activation threshold was applied, To account for any gross scaling differences in effect sizes across contrasts and sessions that could affect the this ICC (McGraw & Wong, 1996), effect sizes were unit normalized within each map prior to each analysis by dividing the value at each vertex by the Euclidian norm of all the vertices in the map.

#### 2.5.3. Vertex-wise Reliability

As in previous fMRI reliability studies (Aron et al., 2006; Caceres et al., 2009; Freyer et al., 2009; Meltzer et al., 2009), we used the ICC to determine the vertex-wise reliability of individuals across sessions. This analysis used the ICC(1) as in 2.5.2, but we defined *MS*_*b*_ in Eq. 4 as the mean squares between subjects, while *MS*_*s*_ and *k* remained the same. Then, to focus our results on vertices that exhibited ‘good’ or ‘excellent’ reliability, we used Koo & Li’s (2016) convention to threshold the resulting ICC map, keeping only those vertices with good or excellent reliability (values greater than or equal to 0.75). Because this measure is calculated with respect to the sample variance, it also provides a measure of discriminability – greater differences between subjects leads to higher values. We applied this analysis to all cortical vertices (without a speech network mask) in order to compare the discriminability of vertices within speech-related areas to those not usually associated with speech. As with the previously described analyses, activation values in each map were unit normalized.

#### 2.5.4. Subject Classifier

Machine-learning tools have recently been applied to MRI data to detect whether subject groups (e.g., patient and control) are discriminable by their neural structure and function (see Sundermann et al., 2014 for a review). Here, we trained a nearest-neighbor subject classifier to identify individual subjects from their functional maps, in order to assess both the reliability and discriminability of *speech* and *null* maps (separately) for individual subjects. First, a session map from the 20 subjects who were scanned twice was set aside as the testing map. A randomly selected single-session activation map from all 75 subjects was then used as the training set (excluding the testing map). The training set data were converted to a set of activation vectors, demeaned, and whitened using the observed between-subjects covariance within the training set (Strang, 1998). The nearest-neighbor classifier then selected the subject within the training set that had the smallest Euclidean distance to the test map. This was repeated for all 40 activation test maps in the dataset (2 maps from each of the 20 subjects with repeated measures) and a percent accuracy score was obtained. This whole procedure was repeated 100 times, each time selecting different sets of random single-session activation maps for training, and the mean accuracy value across these repetitions was taken as the classifier predictive accuracy. Bias-corrected and accelerated (BCa) bootstrapping 95% confidence intervals (Efron, 1987) for accuracy were estimated with 1000 resamples.

For this analysis, we used maps that were masked, thresholded, and unit normalized (see Figure 2B for examples). This meant that subjects were classified by the patterns of relative activation within the most active vertices. We also ran this same classifier on *random* maps (described in section 2.4) to provide an estimate of the accuracy expected based on chance, given the thresholding steps and type of classifier used.

### 2.6. Group-level Statistical Analyses

Dice coefficient and single-subject ICC reliability measures from the *speech, null, and random* maps, which were not assumed to follow a normal distribution, were compared using Wilcoxon Signed-Ranks tests. For the single-subject ICC analysis, we also compared individual ICC values with the between-subjects ICC group measure. In addition, we calculated the Spearman correlations between the *speech* and *null* maps in these measures to determine whether reliability in these two conditions was related (i.e. whether high reliability in the *speech* condition corresponded with high reliability in the *null* condition).

### 2.7. Data and Code Sharing Statement

All anonymized data and analysis code are available upon reasonable request in accordance with the requirements of the institute, the funding body, and the institutional ethics board.

## 3. Results

### 3.1. Single-subject Spatial Overlap

The Dice coefficient for each subject’s thresholded *speech* maps compared between scanning sessions can be found in Figure 3A. On average, their Dice coefficient was 0.693 (SD: 0.089), demonstrating approximately 69% spatial overlap of individual activation maps. For individual *null* maps, the Dice coefficient between sessions 1 and 2 are also shown in Figure 3A. On average, individuals had a Dice coefficient of 0.726 (SD: 0.110), indicating about 73% spatial overlap across sessions. To understand how these values would compare to subjects with completely uncorrelated activation maps, *random* maps yielded a Dice coefficient of 0.205 (SD: 0.016; this is expected, since only voxels with the highest 20% of effect sizes in each map were included). For the group comparison, although *speech* scores were lower than *null* scores, this comparison was not significant (z=-1.31, p=0.191). However, both conditions were significantly different from the *random* maps (z = 3.92, p < 0.001 for both). Further, there was no correlation between Dice coefficients for *speech* and *null* maps (Spearman’s r = 0.098, p = 0.681).

**Figure 3.**
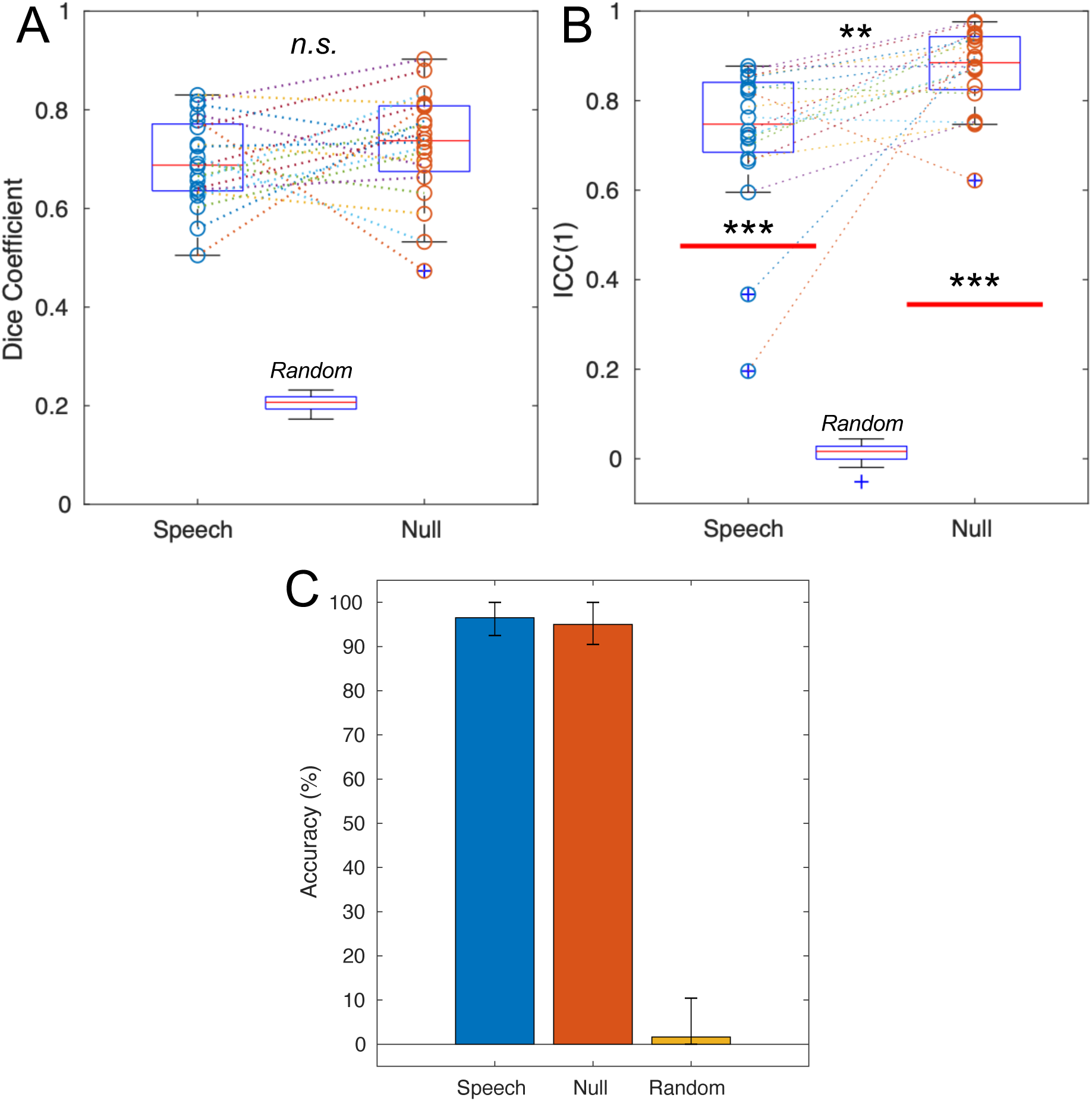
Comparison of reliability measures across conditions. A. Dice coefficient values. Values for individual subjects are shown as circles in each condition, and dashed lines connect results from individual subjects across conditions. For each condition: thin red line = median; blue box = interquartile range (25^th^-75^th^ percentile); black lines = boundary of values for data points that fall within 1.5 times the IQR away from the edges of the box; blue crosses signify outliers – values that fall outside the black lines. B. Single-subject intraclass correlation coefficients. Circles and box plots represent the same information as in A. The thick red lines show the between-subjects intraclass correlation values. Asterisks in line with each condition show comparisons between the distribution of individual points and the Between-Subjects ICC. C. Classifier accuracy. Error bars denote the bias-corrected and accelerated bootstrapping 95% confidence intervals (see section 2.5.4 for details). *n.s*.: non-significant at alpha = 0.05; **: p < 0.01; ***: p < 0.001.

### 3.2. Single-subject ICC

The distribution of single-subject *speech* ICC values across sessions can be found in Figure 3B. Subjects exhibited poor (0.196) to good (0.868) reliability according to the convention of Koo & Li (2016), with a mean ICC(1) of 0.721 (SD: 0.172). As a comparison, the between-subjects correlation, calculated on the averaged individual activation maps across both sessions, was poor with a value of 0.475. A Wilcoxon Signed-Rank test shows that the median of the within-subject ICCs was significantly higher than the between-subject ICC (z=3.51, p<0.001). For the *null* condition, individuals showed moderate (0.622) to excellent (0.976) within-subject reliability, with a mean ICC(1) of 0.870 (SD: 0.092). The between-subjects correlation for this condition was poor at 0.345, and the median of the within-subject coefficients was significantly greater than this value (z=3.92, p<0.001). The *random* maps yielded a mean ICC of 0.013 (SD: 0.025). Within-subject ICCs for the *null* maps were significantly greater than the ICCs for the *speech* maps (z=3.17, p=0.002), and both were significantly greater than random maps (z = 3.92, p < 0.001 for both). Similar to the Dice coefficient, there was no significant correlation between ICC values in the *speech* and *null* conditions (Spearman’s r = 0.173, p = 0.464).

### 3.3. Vertex-wise Reliability

The vertex-wise ICC map for the *speech* data thresholded at 0.75 can be found in Figure 4. While much of cortex was found to have ICC values greater than 0.5 (see Supplementary Figures 1 and 2 for an unthresholded ICC map of *speech* and *null* data), the highest within-subject reliability (>0.75, reflecting good or excellent reliability; Koo & Li, 2016) appeared in areas commonly activated during speech production including, on the lateral surface: bilateral motor and somatosensory cortex, bilateral secondary auditory cortex, bilateral inferior frontal gyrus (IFG) *pars opercularis*, left anterior insula, and bilateral anterior supramarginal gyrus, and on the medial surface: bilateral supplementary and pre-supplementary motor areas, and bilateral cingulate motor area. Some additional regions showed high discriminability as well: bilateral IFG pars orbitalis, right anterior insula, bilateral middle temporal gyrus, and bilateral posterior cingulate cortex. Thus, the speech production network accounts for most of the regions with high within-subject reliability.

**Figure 4.**
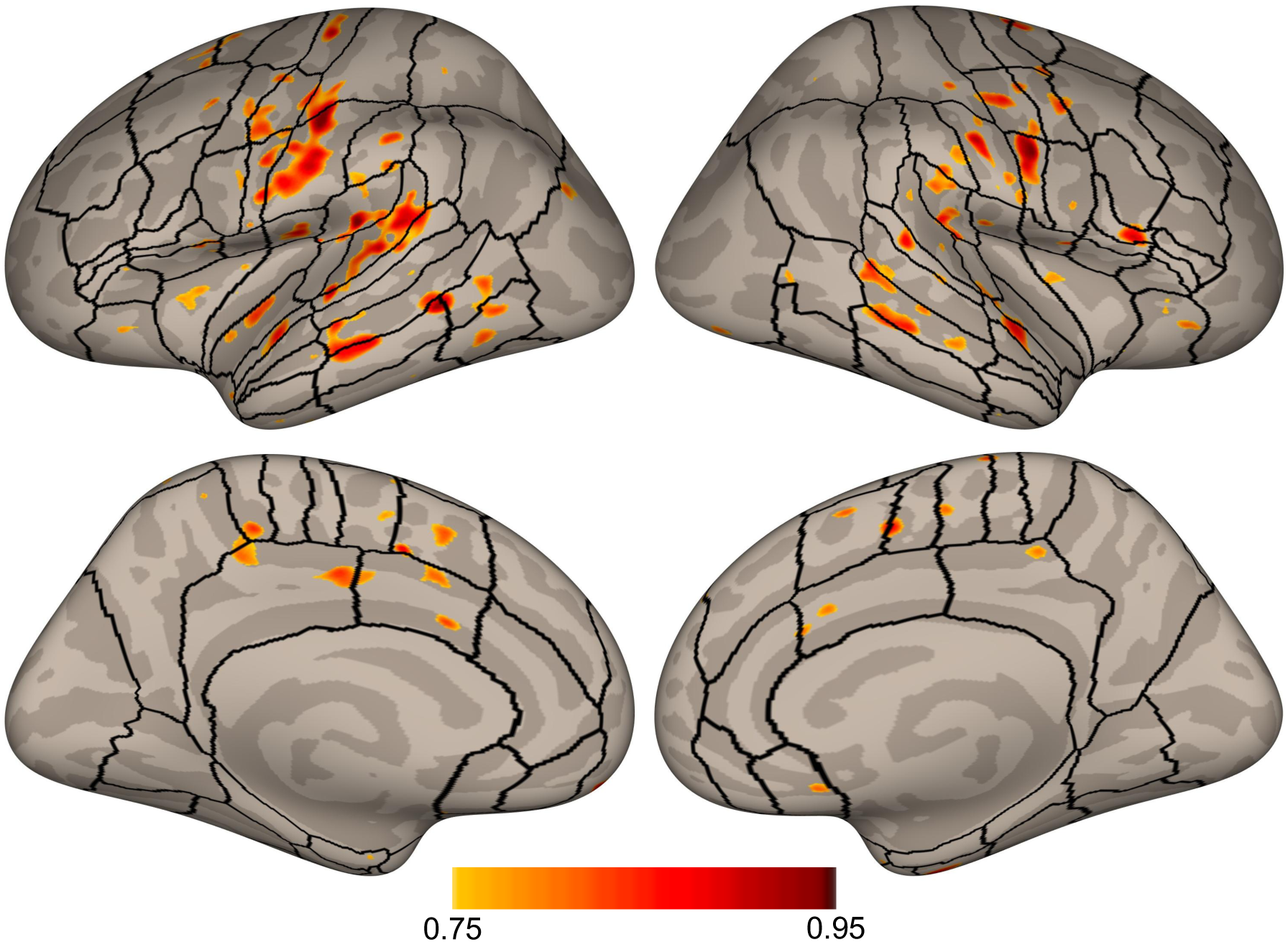
Vertex-wise ICC values for the *speech* activation maps thresholded at 0.75. Regions of interest previously described in Tourville & Guenther (2012) are included for reference.

### 3.4. Subject Classifier

Accuracy of the subject classifier for the *speech* and *null* maps is displayed in Figure 3C. For the *speech* maps, classifier accuracy for untrained test data was 96.52% (BCa bootstrapping 95% confidence interval: 92.5% – 100%). Similarly, the accuracy of this classification method reached 95% for the *null* activation maps (BCa bootstrapping 95% confidence interval: 90.48% – 100%). To assess whether these results were better than chance, we substituted *random* maps for each subject’s *speech* surface maps (while maintaining the number of maps that each subject has and the thresholding pipeline). These results show that for random data, the classifier accuracy was 1.63% (BCa bootstrapping 95% confidence interval: 0% – 10.42%).

## 4. Discussion

Characterizing individual reliability in speech activation is an important step toward validating subject-specific speech research in persons with and without speech disorders. In this study, we used four methods to assess reliability in a group of 20 healthy speakers.

### 4.1. Subject-specific Reliability

The Dice coefficient and single-subject ICC results in this study demonstrated that both the extent and degree of activation patterns during speech production in most, but not all, individuals showed moderate to high amounts of reliability across tasks and timepoints. The Dice values found in this study were generally larger than those found in previous overt expressive language studies (Gorgolewski et al., 2013; Nettekoven et al., 2018; Paek et al., 2019; Wilson et al., 2017). There are several possibilities as to why this was the case. First, the high number of trials for each subject included herein likely increased power which could have improved the robustness of the activation patterns. Indeed, in Gorgolewski et al. (2013), participants had 36 speech trials and 36 baseline compared to an average of 271.9 speech trials in the present analysis (range: 143 – 378) and 78.7 baseline trials (range 24 – 126), likely leading to differences in power as shown previously (Friedman et al., 2008). Paek et al. (2019), on the other hand, included 60 speech trials and 60 baseline trials which may have contributed to its relatively higher Dice coefficients. It is important to note that sessions in these studies were intentionally shortened to accommodate clinical populations, future studies will need to determine how to balance the duel needs of maximal power with minimal scan time.

In addition, as previously discussed, the Dice coefficient is inherently tied to the thresholding scheme used. Gorgolewski et al. (2013), Nettekoven et al., (2018), and Paek et al. (2019) used statistically thresholded maps rather than effect size maps with a percent threshold. Statistically thresholded maps such as these can be strongly affected by multiple factors including noise from head motion and total scan time (Bennett & Miller, 2010; Gross & Binder, 2014). Furthermore, even at similar levels of thresholding (Wilson et al., 2017), reducing the region of interest to pre-defined cortical speech areas in the present study eliminates extraneous regions that show session-specific activations not related to speech *per se*. In Wilson et al. (2017), Dice values in predefined language regions were notably lower than when they looked at all supratentorial voxels, suggesting that higher-level language processing may lead to more variable activation, have lower signal change, and/or contain more noise. Gorgolewski et al. (2013) reported the opposite effect, although Dice values for this task were only specified for auditory cortices. Finally, the older cohorts used in Gorgolewski et al. (2013; age range: 50-58 years), Wilson et al. (2017; age range: 70-76 years), and Paek et al., (2019; age range: 64-83 years) may have had reduced reliability due to various factors that decrease signal-to-noise ratio in the BOLD signal in older adults (D’Esposito et al., 2003). Future work will have to confirm the relationship between age and speech activation reliability.

The single-subject ICC applied in this study measured the degree of reliability between two cortical activation maps. While it relied only on within-subject sources of variance, it was highly correlated with the Dice coefficient (*speech*: Spearman’s r = 0.902, p < 0.001; *null*: r = 0.949, p < 0.001) thus demonstrating its validity as a measure of reliability. One noteworthy difference between this measure and the Dice coefficient was significantly higher ICC for the *null* maps compared to that of the *speech* maps with some subjects attaining near perfect between-session *null* map correspondence. This demonstrates that once all task and motion parameters are accounted for, the underlying signal patterns that reflect individual anatomy maintain high reliability for individuals across scanning sessions. Nonetheless, both *speech* and *null* maps generally demonstrated greater within-subject reliability than a matched between-subjects measure.

There were, however, two participants (Subject 6 and Subject 7) whose within-subjects ICC scores for the *speech* maps were less than the between-subjects ICC estimate. In both cases, the median beta value across vertices for one of the two scanning sessions (the CCRS study session) was more negative than that of any other subjects. This might imply that these subjects had less power for the *speech* contrasts in CCRS. Although they had similar numbers of speech trials as the other subjects, they were among the subjects with the highest scan-to-scan motion and global signal change for this study. They also had the two highest scan-to-scan global signal change values for the other study (FRS). Changes in global signal are often artifacts associated with subject motion, (although other physiological sources contribute to this measure; see Liu, Nalci, & Falahpour, 2017) which was found to be detrimental to reliability measures in previous work (Gorgolewski et al., 2013). However, their motion was not excessive for typical neuroimaging sessions and other subjects with similar amounts of scan-to-scan motion and signal change maintained among the highest ICC values. Another potential reason that these two subjects had much lower ICC scores is methodological: since the ICC(1) measures absolute agreement rather than consistency (McGraw & Wong, 1996), it does not account for global differences in effect sizes across studies. Indeed, the distribution of activation values was shifted between the two sessions to a greater extent for these subjects than for others. We attempted to correct for this by unit-normalizing vertex values for each subject in each study, but this is not a perfect method. Thus, both data quality and methodological choices likely drove down their reliability scores. Minimizing motion will therefore be especially important for future subject-specific analyses.

In sum, we found high within-subject reliability of activation in the speech network, except in two cases where motion may have negatively impacted the signal-to-noise ratio.

### 4.2. Population-normed Reliability

The other two measures we calculated assessed population-normed reliability by comparing response variability within subjects (across sessions) to variability between subjects. These measures assess individual reliability relative to the sample, but additionally characterize how discriminable individuals are from one another. The vertex-wise *speech* ICC map paralleled previous studies that calculated this metric – many of the areas where ICC values were high corresponded to areas commonly activated during the task (Aron et al., 2006; Caceres et al., 2009; Freyer et al., 2009; Meltzer et al., 2009). Thus, for speech production, speech-related areas in somato-motor cortex, medial and lateral pre-motor cortex and extended areas of auditory cortex were consistent for individual subjects across scanning sessions. In addition, even areas of cortex inconsistently active during speech production like IFG *pars orbitalis*, middle temporal gyrus (MTG), and posterior cingulate gyrus (PCG) showed high discriminability. In a review of fMRI studies of speech and language processing (Price, 2012), both IFG pars orbitalis and MTG were associated with semantic processing, while MTG was also associated with translating orthography into sound. This second explanation would be relevant because all tasks involve reading aloud, but it is less clear why semantic processing centers would be highly reliable for pseudoword speaking tasks. The PCG is part of the default mode network and appears to help modulate attentional control (Leech & Sharp, 2014). Thus, individuals may consistently activate or deactivate this region depending on their level of attention during speaking tasks. Previous studies of higher-level cognitive tasks have found reliable activation outside of areas commonly associated with the task, but this usually occurred in sensory and motor regions needed to complete the task (Aron et al., 2006; Freyer et al., 2009). Caceres et al. (2009) suggested that areas with high reliability but low significance values have time-series that are reliable but do not fit the task/HRF model, and demonstrated this pattern for half of their participants in one ROI. This may also be the case in the present study.

It may be worth pointing out that bilateral primary auditory cortex appears less reliable by this vertex-wise ICC measure. While it is counter-intuitive that a low-level sensory region of cortex would be least reliable, this may be an example of one of the drawbacks of this type of measure – since between-subject variance is an important component of this calculation, areas that are more reliable *across* speakers would tend to have *lower* ICC values, given constant within-subject reliability. Thus, it may be more accurate to say that vertices with a high ICC value in this map are the most discriminable areas among a group of subjects.

The final measure of population-normed reliability was the classifier analysis. This type of analysis, which has not previously been used to determine the reliability of an individual’s neural activation patterns, has the added advantage of characterizing the distinctiveness of an individual’s brain activation maps. From the near perfect accuracy in identifying a subject correctly from among 75 potential classes given 1 training sample, it is clear that individuals are not only quite reliable but also have distinct activation patterns during speech production akin to a neural “fingerprint.” In fact, the only subject that was ever mis-classified was Subject 7, who also had the lowest within-subject ICC value and Dice coefficient, thus demonstrating consistency across measures. The same classification method trained on the *null* maps also demonstrated high accuracy, roughly equivalent to that achieved by the *speech* map classifier. It is important to mention that the classification method used in the current study is among the simplest of modern machine learning options, and that using only one training map per subject severely reduces the power of the method. Nonetheless, classification accuracy was very high. We thus interpret the current result as a lower bound of discriminability of speech activation maps among individuals which might be improved with more sophisticated machine learning algorithms.

### 4.3. *Speech* vs. *Null* Reliability

As expected, the portion of the mean BOLD signal associated with brain morphology and neurovasculature demonstrated high reliability within subjects and high discriminability. However, the lack of a correlation between reliability measures in the *speech* and *null* maps suggests that unique activation patterns during the speech task are not dependent on underlying individual anatomy.

### 4.4. Reliability for Speech Production across Tasks

The speech tasks used to assess within-subject reliability herein differed across sessions. This has two important consequences for interpretation of the results. First, the present results do not account for activation variance attributable to inter-task reliability. There may be differences in activation between the studies simply because the speech stimuli were different. Thus, they are potentially conservative compared to the results for a consistent speaking task as well as other published fMRI reliability literature. Second, it means that the reported reliability (and discriminability) measures reflect consistency of the speech production network response rather than the response to a particular task. Therefore, the results are more generalizable to other speech production tasks (at least of the same characteristics – reading orthographic representations of mono- and bi-syllabic words and pseudowords). This is important for assessing the validity of future subject-specific analyses that use speaking tasks that depart from those in the present study.

## 5. Conclusion

Based on the results of four measures of reliability, we conclude that speech activation maps for most neurologically-healthy speakers are generally highly reliable, providing justification for subject-specific neuroimaging research of speech production. Exceptions were found for subjects who exhibited higher levels of scan-to-scan motion and signal change, reinforcing the widely-held understanding that minimizing motion is crucial for trusting neuroimaging data. Future work analyzing activation patterns from patients with neurogenic speech disorders will be needed to determine whether these individuals are similarly reliable (though extant work examining reliability in patients with stroke [Kimberley et al., 2008] and mild cognitive impairment [Zanto et al., 2014] are promising), and ultimately whether subject-specific neuroimaging techniques can be used to map the speech production network in individuals and track changes in these patterns across time. This future research would be an important contribution to the growing body of literature characterizing disease progression and neurorehabilitation (Herbet et al., 2016; Reinkensmeyer et al., 2016), and has the potential to improve diagnosis and treatment for people with speech disorders.

## Acknowledgements

This research was supported by the National Institutes of Health [R01 DC002852, R01 DC007683, and T32 DC013017]. Imaging data were acquired at the Athinoula A. Martinos Center for Biomedical Imaging using resources provided by the Center for Functional Neuroimaging Technologies, P41RR14075 a P41 Regional Resource supported by the National Institute of Biomedical Imaging and Bioengineering, NIH.

## Footnotes

1 Derived from a search of articles on pubmed.com on February 25, 2020 containing the terms “fMRI” or “functional magnetic resonance imaging” and “speech” or “language” in their title or abstract.

