## Supplementary Figures for "Reliability of single-subject neural activation patterns in speech production tasks"

Supplementary Materials

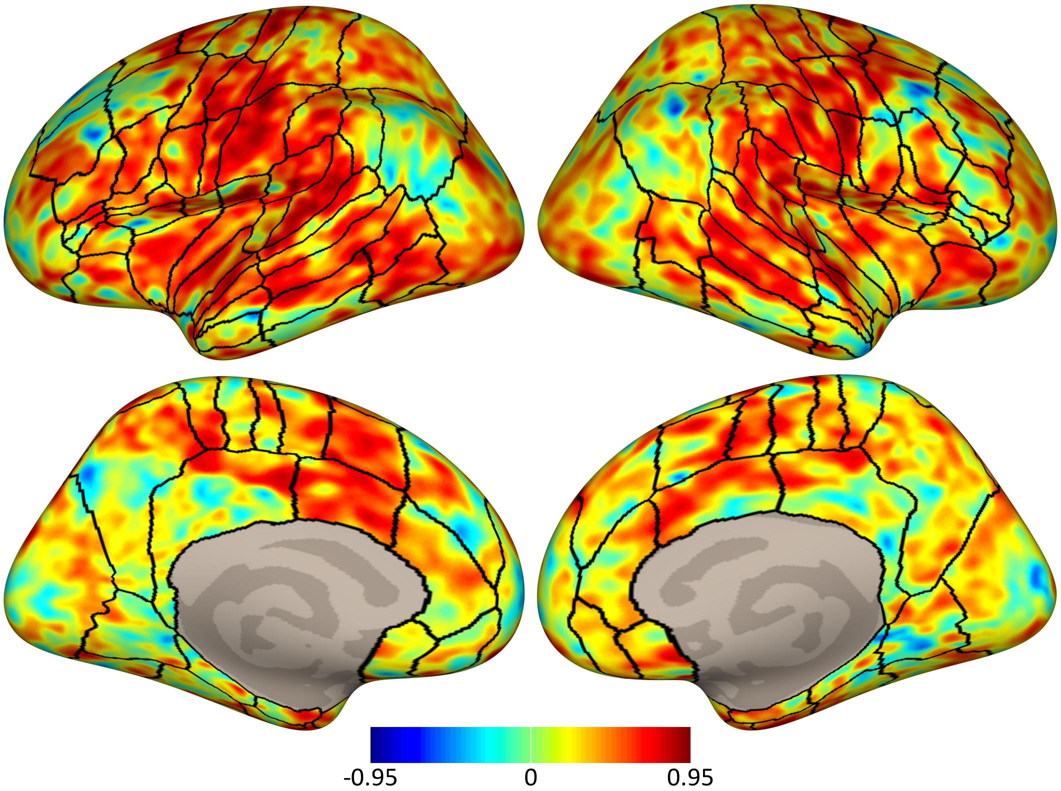


Supplementary Figure 1. Unthresholded vertex-wise ICC values for the *speech* activation maps. Regions of interest previously described in Tourville & Guenther (2012) are included for reference.

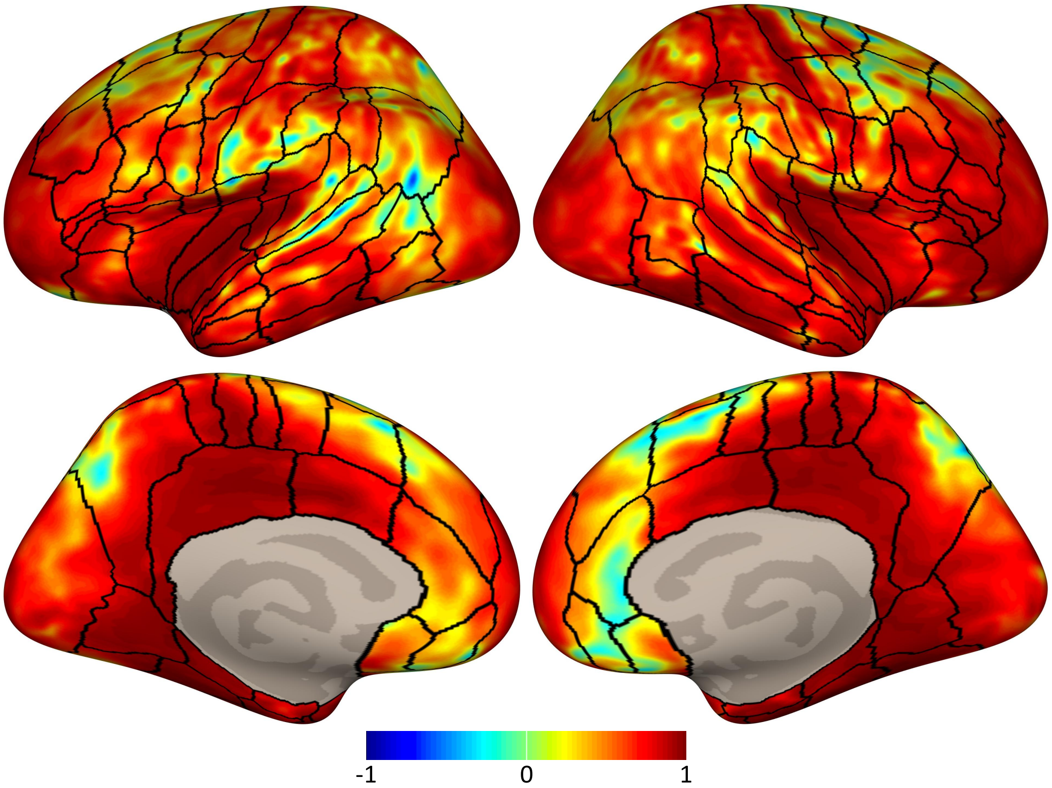


Supplementary Figure 2. Unthresholded vertex-wise ICC values for the *null* activation maps. Regions of interest previously described in Tourville & Guenther (2012) are included for reference.
